# AREION: A Foundation Model for Atlas-Agnostic Representation Learning of Brain Connectome

**DOI:** 10.64898/2026.09.26.754551

**Authors:** Zhao Zhang, Hongming Li, Yuanwang Zhang, Guoliang You, Cong Wang, Yong Fan

**Affiliations:** Department of Electrical and Systems Engineering, University of Pennsylvania, Philadelphia PA, USA; Department of Bioengineering, University of Pennsylvania, Philadelphia PA, USA; Department of Radiology, Perelman School of Medicine, University of Pennsylvania, Philadelphia PA, USA

## Abstract

Brain foundation models have shown promising advances in learning generalizable representations from functional magnetic resonance imaging (fMRI) data. However, existing models are typically tied to specific brain atlases and cannot be applied to fMRI data represented using different atlases without full retraining, thereby severely limiting their broader utility across neuroimaging studies. To address this bottleneck, we introduce *AREION*, **a**tlas-agnostic **re**presentat**i**on learning of brain c**on**nectomes. AREION overcomes atlas dependency through two integrated innovations. First, a *Brain Region Contextualizing* technique maps brain regions and connectivity entries from diverse atlases into a unified representation space based on their spatial, functional, and morphological characteristics. Second, a *Hierarchical Brain Transformer Encoder* processes region-specific connectivity profiles as variable-length token sequences and jointly captures local functional structure and global brain functional organization, conditioned on brain regional context, without assuming a fixed number of regions. Together, these two techniques enable the learning of generalizable representations across brain atlases with different parcellation resolutions and connectivity dimensions. AREION is pretrained on a large-scale multi-atlas cohort of fMRI data via self-supervised modeling and evaluated across diverse downstream tasks. Extensive experiments show that it outperforms strong supervised baselines and fMRI foundation models while using substantially fewer parameters, demonstrating strong transferability to atlases unseen during pretraining, generalization to external datasets, and excellent interpretability at the brain regional level.

## 1 Introduction

Understanding the human brain is of fundamental importance to both neuroscience and clinical applications (Fornito et al., 2015; Drysdale et al., 2017). As a widely used tool in neuroscience, functional magnetic resonance imaging (fMRI) measures brain activity by capturing blood-oxygen-level-dependent (BOLD) signals. Driven by recent advances in artificial intelligence, advanced machine learning approaches have been developed for fMRI analysis (Kawahara et al., 2017; Li et al., 2021; Kan et al., 2022; Chen et al., 2026), leading to improved disease diagnosis and deeper insights into brain function organization. Despite this promise, machine learning models for fMRI analysis are often constrained by limited availability of labeled data and task-specific training paradigms, which restrict their transferability to unseen tasks and datasets. Foundation models offer a promising alternative by pretraining on large-scale unlabeled data to learn generalizable representations that can be adapted to diverse tasks, motivating a growing effort to build foundation models for neuroimaging data.

Recent studies have developed fMRI foundation models for both time-series (Ortega Caro et al., 2024; Dong et al., 2024) and functional connectivity (FC) (Yang et al., 2024; Hu et al., 2024) data. While time-series-based models learn representations directly from the temporal dynamics of fMRI signals, FC-based approaches leverage precomputed connectivity matrices to provide a compact characterization of functional interactions among brain regions, facilitating large-scale pretraining. However, the representations learned by existing FC-based models are tied to the brain atlas used. Such atlas dependence is a fundamental limitation because studies routinely differ in the atlases they adopt, and training a separate model for each parcellation is neither scalable nor efficient. A recent graph-based approach (Wei et al., 2026) takes a step toward addressing this limitation by pretraining across multiple brain atlases and incorporating atlas names as language prompt tokens to facilitate cross-atlas adaptation. However, its ability to generalize to fMRI data represented using unseen atlases from external datasets remains unexplored.

The key challenge in achieving atlas-agnostic representation learning is to construct representations that are transferable across distinct atlases, including those unseen during pretraining. This challenge arises from two related limitations of existing approaches. First, previous work uses textual descriptions of atlas names to derive language embeddings (Wei et al., 2026). However, these embeddings do not explicitly characterize the intrinsic spatial, functional, and morphological properties of individual brain regions of different atlases, particularly those unseen during pretraining. Building an atlas-agnostic fMRI foundation model therefore requires a more effective mechanism to encode regional properties into the learned representations. Second, existing methods typically encode the entire connectivity profile of each region as a single token (Yang et al., 2024; Hu et al., 2024). Because the model dimension depends on the number of regions defined by the adopted atlas, such a representation is tied to a specific parcellation and hinders transfer across atlases with different numbers of regions. Moreover, projecting an entire connectivity profile into a single token overlooks the rich structural dependencies among individual functional connections, limiting model interpretability. Figure 1 summarizes these limitations and the motivation for our framework.

**Figure 1:**
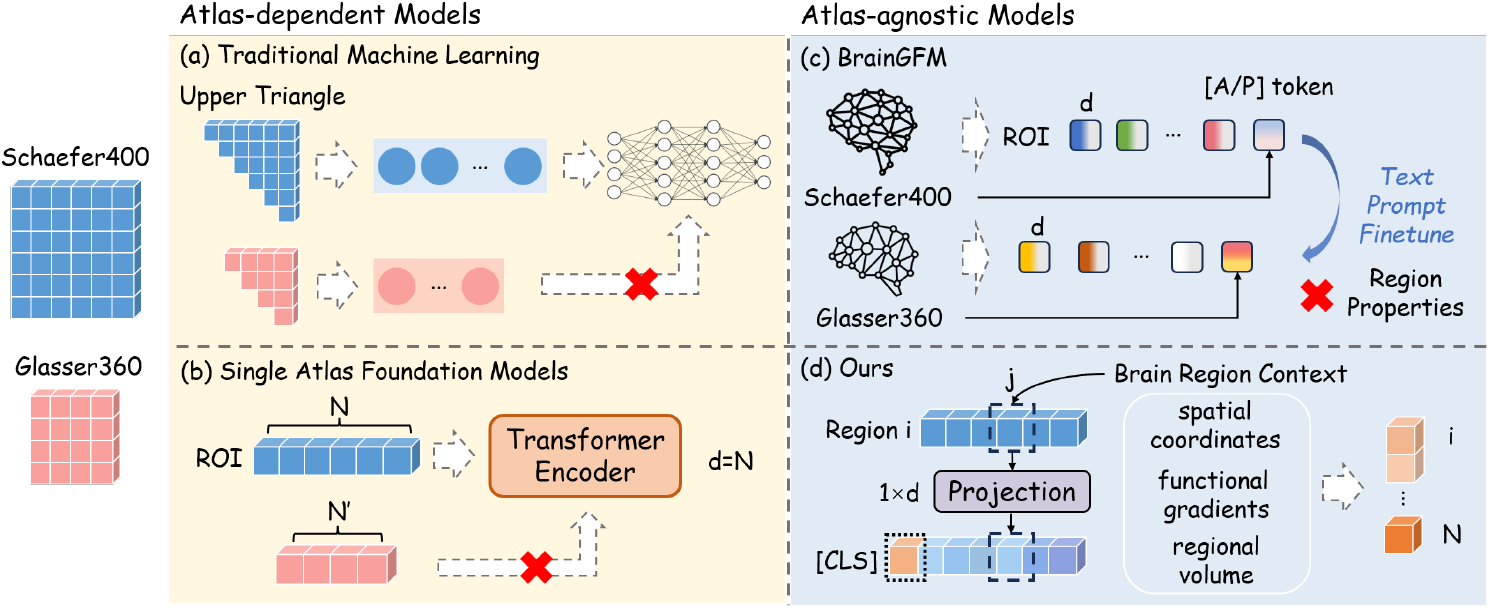
Motivation. Given inputs from distinct brain atlases, **(a)** traditional machine learning methods represent FC matrices as flattened upper triangular vectors, whose dimensions vary across atlases; **(b)** single-atlas foundation models set the model dimension to the number of regions, requiring an atlas-specific configuration; **(c)** BrainGFM enables cross-atlas adaptation by introducing language prompt tokens of atlas names but does not explicitly model the intrinsic characteristics of individual atlases; **(d)** our model maps regional connectivity profiles into a unified representation space by incorporating the spatial, functional, and morphological properties of individual brain regions, thereby enabling cross-atlas transfer.

To address this challenge, we propose AREION, an atlas-agnostic representation learning approach, and use it to build an fMRI foundation model pretrained on 26,000 resting-state fMRI samples from 11,189 unique subjects across five brain atlases. AREION incorporates two key innovations. First, we design Brain Region Contextualizing, a mechanism that encodes continuous information about each brain region by mapping its spatial, functional, and morphological metadata into a shared latent space, allowing the model to adapt to unseen parcellations at inference time without retraining. Second, we design a hierarchical encoder architecture that applies self-attention over variable-length region sequences of functional connectivity profiles rather than operating on a fixed number of regions. This design enables model training on data across multiple atlases while capturing both region-specific connectivity patterns and global brain organization. We conduct extensive experiments comparing AREION with traditional machine learning methods, supervised deep learning models, and pretrained foundation models across diverse downstream tasks, including disease and sex classification and the prediction of age and behavioral measures. Our contributions can be summarized as follows:

- We introduce Brain Region Contextualizing (BRC), a mechanism that represents brain regions using their spatial coordinates, functional gradients, and morphological properties. Unlike previous approaches that rely on atlas names to encode brain regions, BRC directly captures the intrinsic properties of individual regions and provides a unified representation space that can be directly used to characterize brain regions from previously unseen atlases.
- We develop a hierarchical Transformer architecture that separately models fine-grained, region-specific connectivity profiles and whole brain inter-regional dependencies. By treating regional connections as variable-length sequences, the architecture accommodates atlases with different numbers of regions.
- AREION outperforms strong traditional machine learning methods, supervised deep learning models, and existing SOTA fMRI foundation models across diverse downstream tasks and datasets. More importantly, it demonstrates strong transferability to brain atlases unseen during pretraining and provides interpretability at the level of individual brain regions.

## 2 Related Work

### 2.1 Brain Functional Network Analysis

Functional connectivity derived from fMRI data, which is typically estimated as the pairwise Pearson correlation between BOLD time series of regions of interest (ROIs), has been widely used as a biomarker for behavioral traits and neurological disorders. In recent years, deep learning methods have been increasingly applied to brain functional network analysis. Early approaches primarily relied on convolutional architectures, such as BrainNetCNN (Kawahara et al., 2017), which introduces specialized filters to capture the inherent structure of brain networks. Beyond CNNs, a growing number of graph neural networks (GNNs) have been developed (Li et al., 2019; Kim & Ye, 2020; Zhang et al., 2022; Chen et al., 2026), which explicitly model brain regions as nodes and their functional interactions as edges. For instance, BrainGNN (Li et al., 2021) proposes an ROI-aware graph convolutional layer to extract features from brain fMRI graphs, together with an ROI-pooling layer that selects the most informative nodes for downstream tasks. More recently, Brain Network Transformer (Kan et al., 2022) employs Transformer encoders to learn node features in brain networks, and proposes an orthonormal clustering readout layer to exploit modular-level similarities between ROIs. Despite these advances, these approaches are primarily designed for task-specific prediction, limiting the transferability of the learned representations across datasets.

### 2.2 Self-Supervised Learning and Masked Modeling

Self-supervised learning (SSL) has emerged as a powerful paradigm for learning robust representations from large amounts of unlabeled data through a variety of pretext tasks (Gidaris et al., 2018; He et al., 2020; Chen et al., 2020; 2021). By learning from the inherent structure of the data rather than from predefined downstream labels, SSL enables models to capture general patterns that can be transferred across tasks. Among these paradigms, masked autoencoder is a form of generative pretraining in which masked content is reconstructed from the remaining visible data. It was popularized in natural language processing by BERT (Devlin et al., 2019), and was later extended to computer vision by Masked Autoencoders (MAE) (He et al., 2022), which demonstrated that reconstructing masked image patches yields highly transferable visual representations. The success of masked modeling in the language and vision domains has motivated its adaptation to modeling of fMRI data, where large-scale unlabeled fMRI recordings provide an ideal setting for self-supervised pretraining.

### 2.3 fMRI Foundation Models

The success of large-scale pretraining has motivated a growing body of work on fMRI foundation models. Among them, FC-based models operate directly on precomputed FC matrices, providing a compact representation of brain functional organization. BrainMass (Yang et al., 2024) learns FC representations with a Transformer encoder using masked reconstruction and latent-alignment objectives, while BrainNPT (Hu et al., 2024) pretrains a Transformer encoder with replaced ROI prediction before the encoder is finetuned for downstream classification. Although both models achieve competitive downstream performance, they are tied to fixed and predefined parcellations, limiting their direct transfer across atlases that differ in the number and definition of regions. More recently, BrainGFM (Wei et al., 2026) introduces a graph-based approach that enables pretraining across multiple brain atlases by incorporating atlas names as language prompt tokens for cross-atlas adaptation. However, the prompt tokens representing atlas names do not explicitly encode the intrinsic spatial, functional, and morphological properties of individual brain regions, motivating the need for a mechanism that maps brain regions from different atlases into a common representation space.

## 3 Method

To enable atlas-agnostic representation learning across different brain atlases, we propose AREION, consisting of four key components: (1) Brain Region Contextualizing to encode the spatial, functional, and morphological properties of individual brain regions; (2) Hierarchical Brain Transformer Encoder to capture both local connectivity patterns and whole brain functional organization; (3) Masked Autoencoding for self-supervised pretraining; and (4) Downstream Adaptation to adapt and evaluate the learned representations on diverse downstream tasks based on fMRI data represented using different atlases, including those unseen during pretraining. An overview of the proposed framework is illustrated in Figure 2.

**Figure 2:**
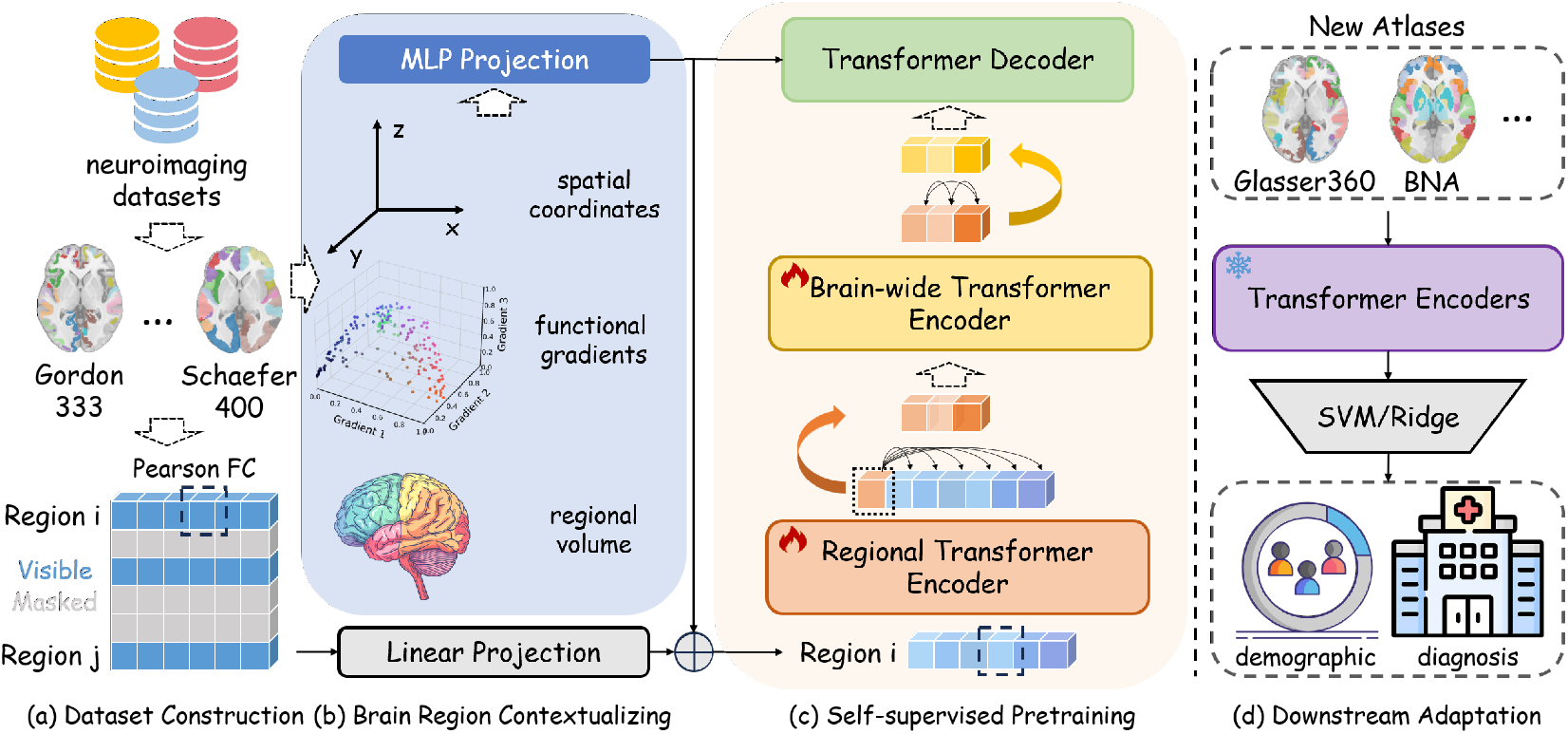
Overview: (a) A large-scale multi-atlas dataset is curated for pretraining; (b) Brain Region Contextualizing encodes the spatial, functional, and morphological properties of individual brain regions; (c) A hierarchical brain Transformer encoder is pretrained via masked autoencoding; (d) The pretrained Transformer encoders are frozen, and the resulting region representations are flattened into a feature vector for downstream prediction.

### 3.1 Definition

Let *A* = {1, …, *N*} denote a brain atlas consisting of *N* ROIs. For a given fMRI scan, we compute the functional connectivity (FC) matrix **F** ∈ ℝ^*N* × *N*^, where each entry *F*_*ij*_ represents the Pearson correlation between the BOLD time series of regions *i* and *j*. Since different atlases yield matrices of different sizes (e.g., *N* = 100, 200, 300, 400, …), we aim to learn an atlas-agnostic encoder 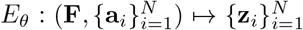 that maps an FC matrix and the associated atlas metadata to a set of regional representations **z**_*i*_ ∈ ℝ^*d*^, one for each region of a given atlas. The encoder is pretrained on unlabeled fMRI data from multiple atlases and subsequently adapted to downstream tasks.

### 3.2 Brain Region Contextualizing

A key challenge in atlas-agnostic representation learning of FC measures is to construct informative representations of brain regional FC profiles adaptive to different atlases. Existing approaches such as BrainGFM incorporate atlas information through language embeddings of atlas or parcellation names. In contrast, we explicitly encode the intrinsic structural and functional properties of each region into a continuous latent space. Specifically, we describe each region *i* with a metadata vector **a**_*i*_ = [**c**_*i*_; **g**_*i*_; *v*_*i*_] ∈ ℝ^13^, comprising its MNI coordinates **c**_*i*_ ∈ ℝ^6^, functional gradient projections **g**_*i*_ ∈ ℝ^6^, and regional volume *v*_*i*_ ∈ ℝ. Through BRC, the spatial, functional, and morphological metadata of an individual region *i* are mapped to a *d*-dimensional latent representation:

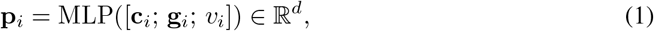

where MLP consists of a linear projection, layer normalization, a GELU activation (Hendrycks & Gimpel, 2016), and a second linear projection with dropout. Here, **p**_*i*_ is computed from the continuous intrinsic properties of the brain region rather than from a fixed region index or an atlas-specific vocabulary, allowing the encoder to handle arbitrary atlases at inference time, and the three inputs of each region capture its intrinsic characteristics independent of atlases: 1) **c**_*i*_ ∈ ℝ^6^: the mean MNI coordinates (*x, y, z*) of the voxels and their corresponding standard deviations (*σ*_*x*_, *σ*_*y*_, *σ*_*z*_), capturing the *spatial* properties of the region; 2) **g**_*i*_ ∈ ℝ^6^: the first three functional gradient projections (*g*_1_, *g*_2_, *g*_3_) and their corresponding standard deviations 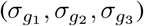, capturing the position of the region along the principal axes of *functional* organization in the brain; and 3) *v*_*i*_ ∈ ℝ: the regional volume, capturing the *morphological* properties of the region. Particularly, the functional gradients are informative for characterizing functional changes associated with age, cognition, and brain disease (Nguyen et al., 2024; Bethlehem et al., 2020). Since each functional connectivity value corresponds to the interaction between two brain regions, the BRC embeddings of regions *i* and *j* are combined additively as **p**_*i*_ + **p**_*j*_ to encode their connectivity so that the properties of both regions are incorporated into their connectivity token.

### 3.3 Hierarchical Brain Transformer Encoders

Motivated by the multi-level organization of human brain networks (Meunier et al., 2009; Sporns & Betzel, 2016), we develop a hierarchical Transformer architecture to separately model regional FC profiles and whole brain FC organization, thereby preserving fine-grained regional level FC patterns while capturing global organization across atlases with varying numbers of brain regions.

#### 3.3.1 Regional Transformer Encoder

The Regional Transformer Encoder (RTE) models the FC profile of each region by operating independently on each row of the FC matrix. For region *i*, its FC profile **F**_*i*_ = [*F*_*i*1_, …, *F*_*iN*_] ∈ ℝ^*N*^ describes its connectivity to all other regions. We project each connectivity value *F*_*ij*_ into a *d*-dimensional token through a learned linear projection combined with the pairwise positional encoding:

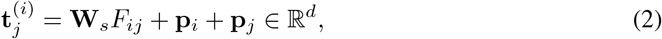

where **W**_*s*_ ∈ ℝ^*d×*1^ is a shared learnable projection. The token 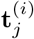 thus encodes both the strength of the connection (*F*_*ij*_) and the properties of its two connected regions. A learnable CLS token **e**_cls_ is then prepended to the sequence to aggregate information across all connectivity tokens through self-attention. The resulting sequence is processed by a Transformer encoder:

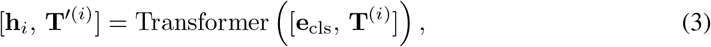

where 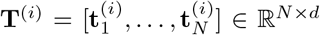. The CLS output **h**_*i*_ ∈ ℝ^*d*^ serves as the region-level representation that summarizes the connectivity profile of region *i*.

#### 3.3.2 Brain-wide Transformer Encoder

The RTE models the FC profile of each region independently but does not explicitly capture whole brain functional organization. To address this issue, we develop a Brain-wide Transformer Encoder (BTE) to jointly model all regional tokens for characterizing the global organization of the functional brain network.

The BTE takes the region-level representations **H** = [**h**_1_, …, **h**_*N*_] as input and adds the corresponding BRC embeddings **P** = [**p**_1_, …, **p**_*N*_] to form enriched input tokens. A deeper Transformer encoder then processes this sequence through global self-attention across all brain regions:

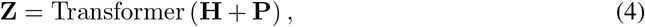

where **Z** = [**z**_1_, …, **z**_*N*_] ∈ ℝ^*N ×d*^ denotes the resulting context-enriched region tokens. These representations capture both region-specific connectivity profiles and whole-brain functional organization and are used as the learned brain representations for downstream tasks.

### 3.4 Masked Autoencoding

We pretrain the Transformer encoders using a masked autoencoding paradigm. Given *N* regions, we randomly sample a visible subset V ⊂{1, …, *N*}, where |V|= (1 −*r*)*N* and *r* denotes the masking ratio. The remaining regions, ℳ = {1, …, *N*} \ V, are masked. The visible regions are then processed by the hierarchical Transformer encoders to obtain regional representations {**z**_*i*_}_*i*∈V_, while the masked regions are excluded from the encoder input. This asymmetric design substantially reduces the computational cost of pretraining because the encoders process only a fraction, 1 *− r*, of the *N* regions.

For reconstruction, each masked region is represented by a learnable mask embedding **e**_mask_ ∈ ℝ^*d*^. The corresponding BRC embedding **p**_*i*_ is added to the learnable embedding so that the decoder retains the spatial, functional, and morphological identity of the region even when the connectivity information is unavailable. The decoder input for region *i* is defined as:

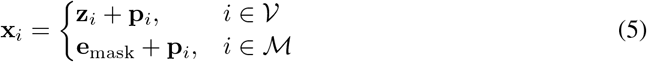

A Transformer decoder processes the complete sequence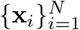 through global self-attention. This allows masked regions to aggregate contextual information from visible region tokens, producing decoded representations 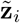 for all *N* regions. Because the FC matrix **F** is computed using Pearson correlations, which correspond to normalized inner products between regional BOLD time series, we reconstruct each element of **F** using a matching inner-product readout:

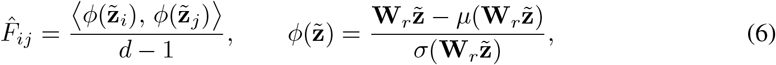

where **W**_*r*_ ∈ ℝ^*d×d*^ is a learnable projection matrix, and *µ*(*·*) and *σ*(*·*) denote the mean and sample standard deviation computed across the *d* channels, respectively. The reconstruction loss is computed over the masked rows of FC matrix:

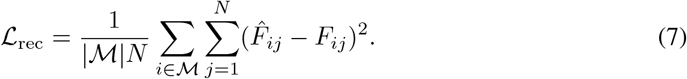

### 3.5 Downstream Adaptation and Applications

For downstream evaluation, we freeze the pretrained Transformer encoders and extract regional representations from FC matrices defined using any atlases. For a given input FC matrix with *N* regions, the encoders produce *N* region tokens of dimension *d*. These tokens are concatenated and flattened into an *N× d* feature vector, which is fed into a task-specific predictor: a support vector machine (SVM) for classification tasks or a ridge regressor for regression tasks. Although regional representations from atlases with varying numbers of regions can be integrated using Transformers, the present study focuses on evaluating each atlas separately using traditional machine learning models that require fixed-dimensional feature vectors.

## 4 Experiments

### 4.1 Datasets and Evaluation

#### Datasets

We pretrain our model on three fMRI datasets: the Human Connectome Project (HCP) (Van Essen et al., 2013), Human Connectome Project-Development (HCP-D) (Somerville et al., 2018), and Adolescent Brain Cognitive Development (ABCD) (Casey et al., 2018), with their fMRI data processed using multiple atlases. For external evaluation, we use the Philadelphia Neurodevelopmental Cohort (PNC) dataset (Satterthwaite et al., 2014), with its fMRI data processed using two atlases unseen during pretraining: Glasser360 atlas (Glasser et al., 2016) and the Brainnetome Atlas (BNA; 210 regions) (Fan et al., 2016). Additional dataset details are provided in Appendix A.

#### Evaluation Metrics and Interpretability Analysis

We evaluate classification performance using accuracy (ACC), area under curve (AUC), and macro-F1 score, and regression performance using the coefficient of determination (*R*^2^), mean absolute error (MAE), and Pearson correlation coefficient (*ρ*). Interpretability analyses of the learned FC representations are provided in Appendix F.

### 4.2 Main Results

#### 4.2.1 Comparison with other Methods Using FC from the Same Atlases

We compare AREION with traditional machine learning (ML) methods, supervised deep learning (DL) models, and pretrained fMRI foundation models. For a fair comparison, all foundation models are retrained on our collected data, and all methods are evaluated using the same five-fold cross-validation. Because BrainGFM and AREION support multi-atlas pretraining, we evaluate both methods under single-atlas and multi-atlas settings. The single-atlas setting is consistent with that used for the other foundation model baselines. Additional details about baselines are provided in Appendix E.

As shown in Table 1, traditional ML methods are highly competitive. SVM and ridge regression outperform several supervised DL and pretrained foundation models across multiple tasks. For example, SVM outperforms BrainGNN and BrainMass on both sex and schizophrenia classification tasks, highlighting the importance of including strong traditional ML baselines. Under single-atlas pretraining, AREION outperforms all foundation models evaluated under the same setting. BrainGFM and AREION also generally benefit from multi-atlas pretraining, with AREION achieving the strongest overall performance across the evaluated downstream tasks. These results support the potential of multi-atlas pretraining to learn more generalizable representations. Additional comparisons across atlas resolutions are provided in Appendix B.

**Table 1:**
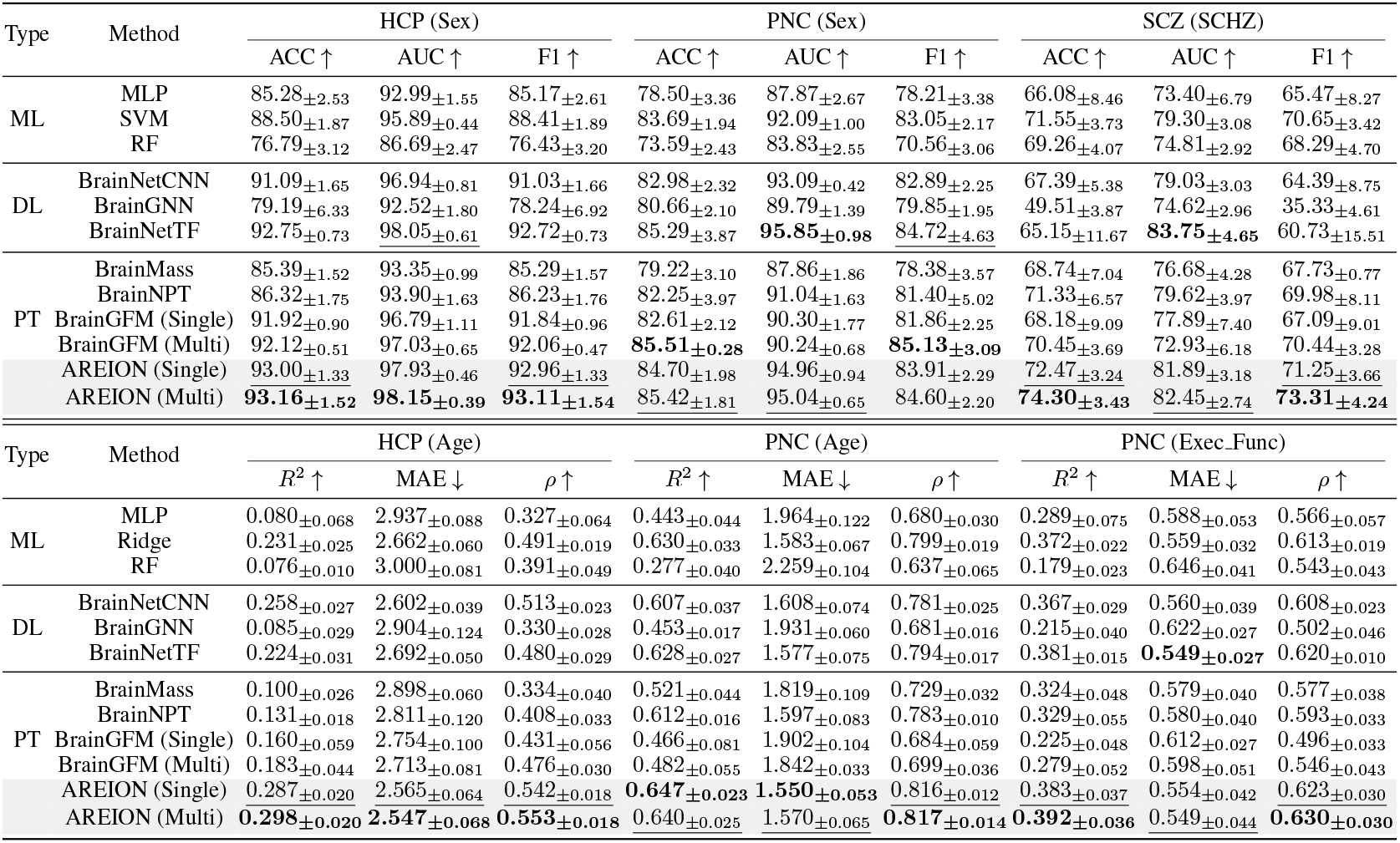
Performance comparison across downstream tasks on the Schaefer400 atlas: sex classification (Sex), schizophrenia classification (SCHZ), age regression (Age), and executive function prediction (Exec Func). The best and second-best results are highlighted in **bold** and underlined, respectively. ML, DL, and PT denote traditional machine learning methods, supervised deep learning models, and pretrained foundation models, respectively. **Note:** BrainGFM (Single) and AREION (Single) are pretrained using only the Schaefer400 atlas, whereas BrainGFM (Multi) and AREION (Multi) are pretrained using multiple atlases.

#### 4.2.2 Generalization to Unseen Atlases

To evaluate AREION’s ability to generalize to unseen atlases, we conduct experiments on the external PNC dataset, with its fMRI data processed using the Glasser360 and Brainnetome atlases, neither of which is used during pretraining. We compare AREION with BrainGFM, which also supports multi-atlas pretraining. Both models are pretrained on our collected cohort. As shown in Table 2, AREION consistently outperforms both the conventional ML baselines and BrainGFM on both unseen atlases, demonstrating strong cross-atlas transferability. In contrast to BrainGFM, which encodes textual descriptions of atlas names, AREION uses Brain Region Contextualizing to directly encode the intrinsic spatial, functional, and morphological properties of individual brain regions. The results support the effectiveness of this regional contextualization in learning representations that transfer reliably to previously unseen atlases.

**Table 2:**
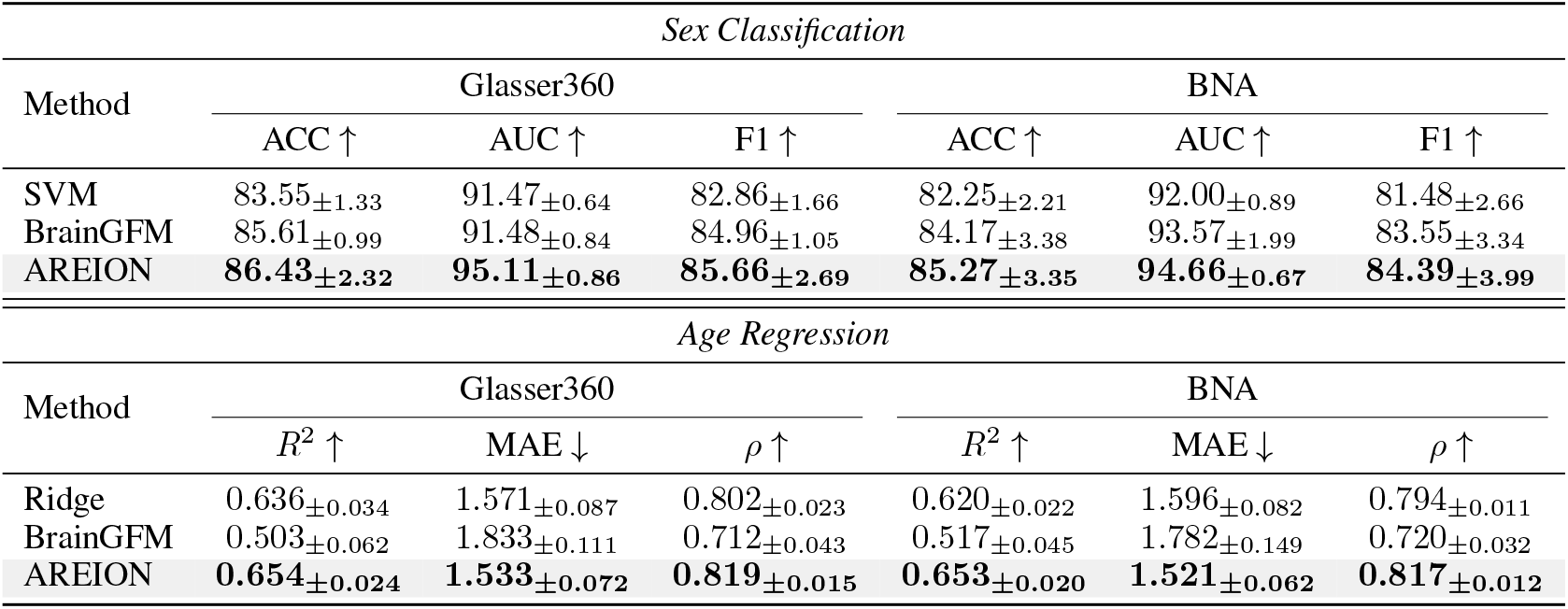
Generalization to unseen brain atlases. We evaluate the generalizability of AREION, SVM, ridge regression, and BrainGFM on the external PNC dataset using atlases unseen during pretraining.

### 4.3 Ablation Studies

#### 4.3.1 Ablation Study of Positional Encoding Methods

To investigate the contribution of BRC to cross-atlas generalization, we compare it with two commonly used positional encoding (PE) methods: sinusoidal positional encoding (Vaswani et al., 2017) and relative positional encoding (Shaw et al., 2018). All variants use the same hierarchical encoder architecture and are independently pretrained on the HCP dataset using the Schaefer100/200/300/400 atlases. We evaluate these variants on FC measures derived from the Schaefer400 atlas, which is seen during pretraining, and the Gordon333 atlas, which is unseen during pretraining.

As shown in Table 3, all positional encoding methods substantially outperform the model without positional encoding, demonstrating the importance of positional information. On the seen Schaefer400 atlas, BRC achieves the best performance across all metrics. Although sinusoidal PE achieves comparable results in this setting, its performance declines notably when transferred to the Gordon333 atlas, whereas BRC maintains more stable performance. These results indicate that, unlike conventional PE methods that rely on absolute or relative sequence positions, BRC encodes continuous spatial, functional, and morphological properties to produce more transferable representations of brain regions, particularly for generalization to unseen atlases.

**Table 3:** Ablation study of different positional encoding methods. All variants are independently pretrained on the HCP dataset and evaluated for sex classification using a seen atlas, Schaefer400, and an unseen atlas, Gordon333, during pretraining.

| Positional Encoding | Schaefer400 |  |  | Gordon333 |  |  |
| --- | --- | --- | --- | --- | --- | --- |
| | ACC ↑ | AUC ↑ | F1 ↑ | ACC ↑ | AUC ↑ | F1 ↑ |
| w/o PE | 75.03±1.20 | 84.02±2.29 | 74.67±1.23 | 72.85±2.63 | 82.04±2.45 | 72.37±2.63 |
| Sinusoidal PE | 91.92±2.16 | 97.48±0.59 | 91.86±2.18 | 88.81±1.72 | 95.17±0.37 | 88.75±1.73 |
| Relative PE | 90.78±0.39 | 96.50±0.61 | 90.72±0.41 | 86.22±1.49 | 94.38±0.47 | 86.13±1.49 |
| **BRC (Ours)** | **92.96±1.24** | **97.91±0.45** | **92.94±1.25** | **91.61±1.20** | **97.17±0.60** | **91.56±1.19** |

#### 4.3.2 Ablation Study on Framework Components

We conduct ablation studies to investigate the contributions of the proposed framework components. All model variants are pretrained on the HCP dataset and evaluated under both in-distribution and cross-atlas settings. We first examine the contributions of the three components of BRC: spatial coordinates, functional gradients, and regional volume. As illustrated in Figure 3(a), incorporating functional gradients substantially improves performance, while the subsequent addition of regional volume consistently yields the highest overall accuracy. These results suggest that the three components provide complementary information about regional identity, enabling the model to generalize more effectively across distinct atlases.

**Figure 3:**
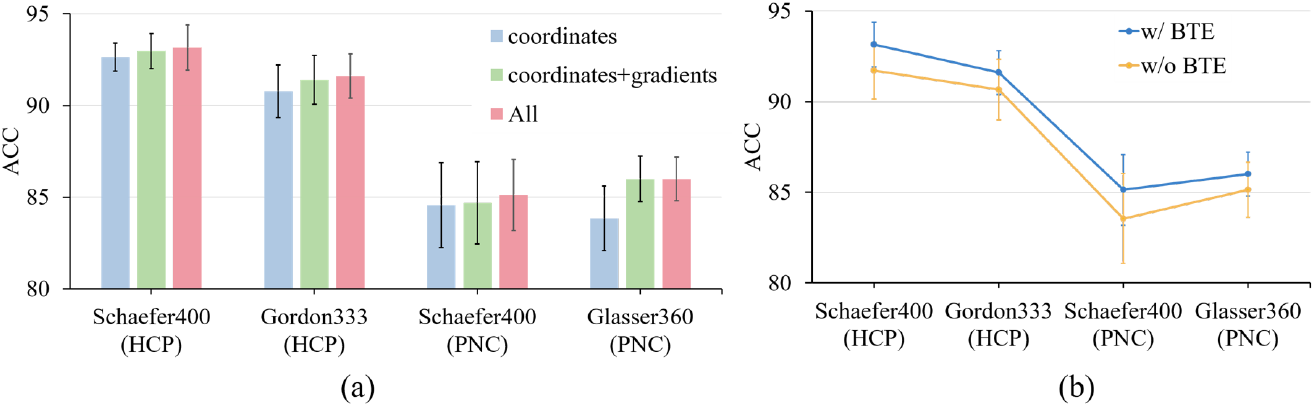
Ablation study of framework components. **(a)** Effects of the spatial coordinates, functional gradients, and regional volume of Brain Region Contextualizing. **(b)** Effect of Brain-wide Transformer Encoder. All models are evaluated on the sex classification task using HCP with the seen Schaefer400 atlas and PNC with the unseen Gordon333 and Glasser360 atlases during pretraining.

Next, we compare AREION with its ablated version in which the Brain-wide Transformer Encoder is removed. As shown in Figure 3(b), the model with the BTE consistently outperforms its counterpart without BTE, demonstrating the effectiveness of the proposed hierarchical representation learning strategy. Without the BTE, the learned representations primarily capture region-specific connectivity patterns. Incorporating the BTE further models inter-regional dependencies and enriches the regional representations with global brain context.

## 5 Conclusion

In this paper, we introduce AREION, an fMRI foundation modeling approach for atlas-agnostic representation learning of brain connectomes. To overcome the dependence of existing approaches on fixed brain atlases, we propose two complementary techniques: Brain Region Contextualizing, which represents brain regions and their connectivity entries using their intrinsic spatial, functional, and morphological characteristics; and a Hierarchical Brain Transformer Encoder, which captures region-specific connectivity patterns and global inter-regional topology from variable-length connectivity sequences. Extensive experiments demonstrate that AREION achieves superior performance across diverse downstream tasks, generalizes effectively to atlases unseen during pretraining, and produces representations with biologically meaningful interpretability at the regional level.

## A Dataset Details and Preprocessing

### Human Connectome Project (HCP)

The HCP dataset contains fMRI scans of healthy subjects aged 22–37 years. Details of the acquisition protocols have been published previously (Glasser et al., 2013). The minimally preprocessed outputs from the HCP pipeline were postprocessed using XCP-D. The processed fMRI scans of 965 subjects were included in this study. For each subject, two resting-state fMRI sessions were included, and two runs with left-to-right and right-to-left phase encoding were concatenated to compute the functional connectivity for each session.

### Human Connectome Project-Development (HCP-D)

The HCP-D dataset includes subjects recruited across four sites in the United States, to study healthy brain development in children and adolescents. The image acquisition and preprocessing procedures have been described previously (Luo et al., 2024). fMRI scans from 632 subjects were included in this study.

### Adolescent Brain Cognitive Development (ABCD)

The ABCD dataset contains fMRI scans from participants aged 9–10 years at baseline, collected across 21 sites in the United States. Resting-state fMRI scans of 9, 592 participants preprocessed by the ABCD BIDS Community Collection (ABCC) (Feczko et al., 2021) were used in this study.

### Philadelphia Neurodevelopmental Cohort (PNC)

The PNC dataset comprises a community-based sample of children and adolescents from the greater Philadelphia area. We included demographic measures, cognitive assessments, and fMRI scans from 693 participants (301 males, 392 females) aged 8–23 years. Subject inclusion criteria and image preprocessing followed previously established protocols (Cui et al., 2020).

### Schizophrenia Dataset (SCZ)

The SCZ dataset consists of 218 individuals, including 117 healthy controls and 101 schizophrenia patients, collected from previous studies (Jing et al., 2019; Zhuo et al., 2017). All subjects underwent structural MRI and resting-state fMRI scanning on a 3.0-Tesla MR system. The resting-state fMRI data were preprocessed using fMRIPrep (Esteban et al., 2019), and motion-related artifacts were reduced using ICA-AROMA (Pruim et al., 2015).

Table 4 provides an overview of the datasets used in our study. We categorize the datasets into four groups according to their roles: pretraining, internal testing, semi-external testing, and external testing. The pretraining group consists of three widely used fMRI datasets, totaling 11,189 unique subjects and 26,000 samples derived using five distinct brain atlases. The internal testing group includes settings in which both the dataset and the atlas are seen during pretraining. It is constructed from subsets of the pretraining datasets and is used to evaluate the in-distribution performance of our approach. The semi-external testing group includes settings in which either the dataset or the atlas is unseen during pretraining, while the other is seen, enabling the evaluation of cross-dataset or cross-atlas generalization. The external testing group includes settings in which both the dataset and atlas are unseen during pretraining, enabling the evaluation of generalization across both unseen datasets and unseen atlases. Collectively, the test groups cover diverse downstream tasks, including demographic prediction, such as age regression and sex classification, cognitive function prediction, such as executive function prediction, and disease diagnosis, such as schizophrenia classification. These tasks enable a comprehensive evaluation of the generalizability of the learned representations.

**Table 4:** Overview of neuroimaging datasets used in this study. We group datasets according to their roles in our pipeline: pretraining, internal testing, semi-external testing, and external testing. (1) **Pretraining**: Data used for self-supervised pretraining. (2) **Internal Testing**: Settings in which both the dataset and atlas are seen during pretraining. (3) **Semi-External Testing**: Settings in which either the dataset or the atlas is unseen during pretraining. (4) **External Testing**: Settings in which both the dataset and atlas are unseen during pretraining. The table lists datasets, atlases, number of unique subjects, total samples, and associated downstream tasks.

| Datasets | Atlases | Subjects | Samples | Task |
| --- | --- | --- | --- | --- |
| *Pretraining* |  |  |  |  |
| HCP | Schaefer100/200/300/400 | 965 | 13,880 | - |
| HCP-D | Schaefer100/200/300/400 | 632 | 2,528 | - |
| ABCD | Gordon333 | 9,592 | 9,592 | - |
| *Internal Test* |  |  |  |  |
| HCP | Schaefer100/200/300/400 | 965 | 3,860 | Multiple |
| *Semi-External Test* |  |  |  |  |
| HCP | Gordon333 | 965 | 965 | Multiple |
| PNC | Schaefer400 | 693 | 693 | Multiple |
| SCZ | Schaefer100/200/300/400 | 218 | 872 | Schizophrenia Classification |
| *External Test* |  |  |  |  |
| PNC | Glasser360/BNA | 693 | 1,386 | Multiple |
| SCZ | Glasser360 | 218 | 218 | Schizophrenia Classification |

## B Additional Results

### B.1 Additional Comparison with Other Baselines

To further examine the effect of parcellation resolution on baseline performance, we compare the methods across multiple brain atlases. Figures 4 and 5 show their performance on the Schaefer100, Schaefer200, Schaefer300 and Schaefer400 atlases for HCP sex classification and age regression, respectively. For BrainGFM and AREION, we report results from models pretrained on multiple atlases, whereas the other methods under comparison are trained or retrained separately for each corresponding atlas resolution. As shown in the figures, AREION maintains consistently strong performance across different parcellation resolutions. Moreover, most baseline methods exhibit improved performance as the parcellation resolution increases, suggesting that higher-resolution parcellations may provide finer-grained regional FC information that benefits downstream prediction.

**Figure 4:**
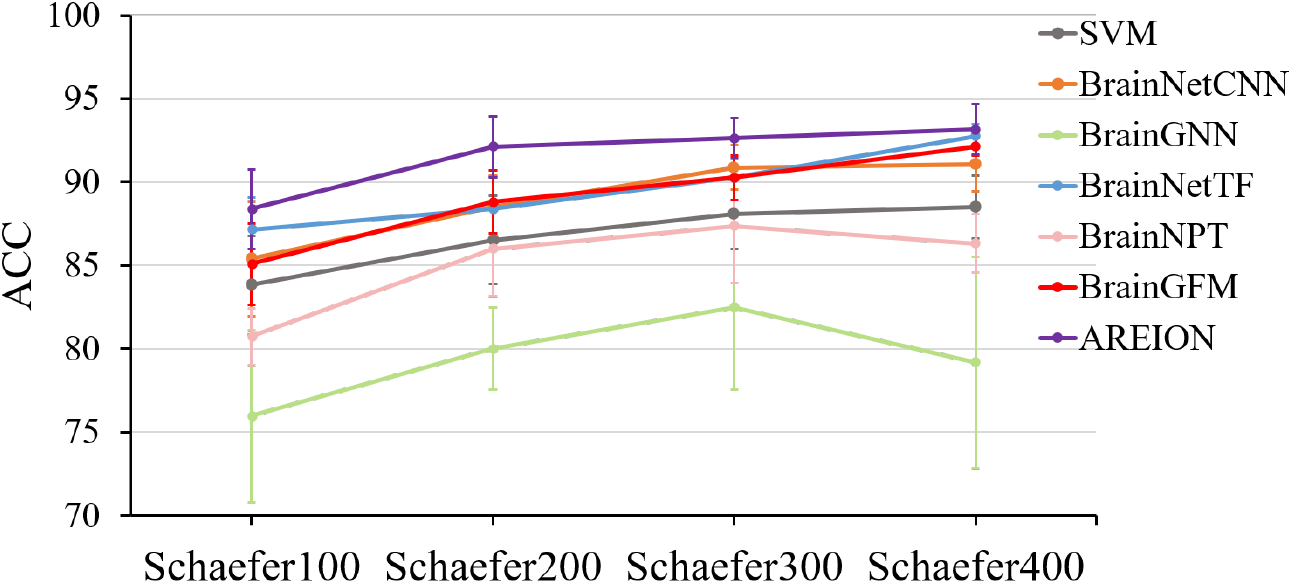
Performance comparison of different methods for HCP sex classification on Schaefer100/200/300/400.

**Figure 5:**
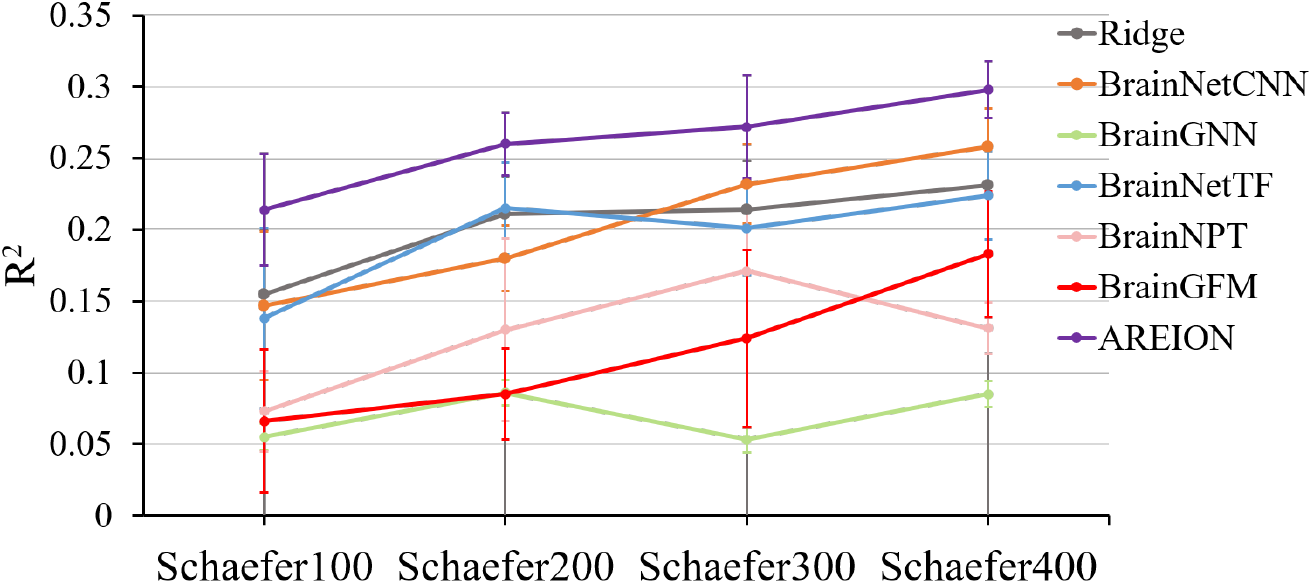
Performance comparison of different methods for HCP age regression on Schaefer100/200/300/400.

### B.2 Additional Ablation Results

To complement the ablation results reported in the main paper, we further evaluate different positional encoding methods on the HCP sex classification task using the Schaefer100, Schaefer200, and Schaefer300 atlases. As shown in Table 5, BRC consistently achieves the best performance across all three parcellation resolutions and all evaluation metrics, further supporting its effectiveness.

**Table 5:** Ablation study of positional encoding methods. All variants are independently pretrained on the HCP dataset and evaluated on the HCP sex classification task using the Schaefer100/200/300 atlases.

| Positional Encoding | Schaefer100 |  |  | Schaefer200 |  |  | Schaefer300 |  |  |
| --- | --- | --- | --- | --- | --- | --- | --- | --- | --- |
| | ACC ↑ | AUC ↑ | F1 ↑ | ACC ↑ | AUC ↑ | F1 ↑ | ACC ↑ | AUC ↑ | F1 ↑ |
| w/o PE | 73.47±1.20 | 80.31±3.09 | 73.15±0.94 | 73.68±1.65 | 81.35±2.72 | 73.32±1.68 | 74.72±1.48 | 82.80±2.67 | 74.33±1.53 |
| Sinusoidal PE | 87.98±2.13 | 95.61±0.61 | 87.90±2.16 | 91.61±2.33 | 96.71±0.72 | 91.56±2.35 | 92.54±0.70 | 97.46±0.42 | 92.51±0.71 |
| Relative PE | 83.52±3.17 | 92.57±1.25 | 83.39±3.23 | 86.74±1.96 | 94.69±0.67 | 86.64±2.02 | 88.70±1.24 | 95.99±0.45 | 88.64±1.28 |
| BRC (Ours) | **88.50±2.35** | **95.77±0.58** | **88.43±2.37** | **91.81±2.16** | **97.20±0.66** | **91.76±2.16** | **92.75±1.47** | **97.73±0.44** | **92.70±1.48** |

## C Implementation Details

For pretraining, we use the AdamW optimizer (Loshchilov & Hutter, 2017) with a learning rate of 1.5 *×* 10^*−*4^ and a weight decay of 0.05. The learning rate undergoes a linear warmup over 40 epochs followed by a cosine decay schedule(Loshchilov & Hutter, 2016). The model is pretrained for 400 epochs with a mask ratio of 75% and a batch size of 8. All pretraining experiments are conducted using two NVIDIA A100 GPUs. Additional pretraining and architectural hyperparameters are provided in Table 6.

**Table 6:** Training and architectural hyperparameters used in the proposed framework.

| Hyperparameter | Value | Description |
| --- | --- | --- |
| batch_size | 8 | Number of samples per batch during training. |
| learning_rate | $1.5 \times 10^{-4}$ | Learning rate used by the optimizer. |
| epochs | 400 | Number of epochs. |
| warmup_epochs | 40 | Number of warmup epochs. |
| weight_decay | 0.05 | Optimizer weight decay coefficient. |
| betas | [0.9, 0.95] | Optimizer momentum. |
| d | 256 | Hidden dimension shared across the Brain Region Contextualizing, hierarchical encoders and MAE decoder. |
| region_nhead | 4 | Number of attention heads in Regional Transformer Encoder. |
| region_layers | 4 | Number of Transformer layers used in the Regional Transformer Encoder. |
| brain_nhead | 8 | Number of attention heads in Brain-wide Transformer Encoder. |
| brain_layers | 6 | Number of Transformer layers used in the Brain-wide Transformer Encoder. |
| mask_ratio | 0.75 | Fraction of regions masked. |
| dropout | 0.1 | Dropout rate used for regularization. |

For downstream adaptation, we freeze the pretrained Transformer encoders and evaluate the output embeddings using SVM for classification tasks and ridge regression for regression tasks. The pretrained encoder is not finetuned for the downstream evaluation. More settings about the architectures of downstream models are provided in Table 7.

**Table 7:** Downstream evaluation hyperparameters used in the proposed framework.

| Hyperparameter | Value | Description |
| --- | --- | --- |
| **SVM** |  |  |
| kernel | RBF | Kernel function used for classification. |
| probability | True | Enables probability estimates for AUC computation. |
| C | 1.0 | SVM regularization strength. |
| gamma | scale | RBF kernel coefficient. |
| **Ridge** |  |  |
| alpha | $\{10^{-1}, \dots, 10^6\}$ | 30 log-spaced candidate L2 regularizations. The best value is selected using leave-one-out cross-validation via RidgeCV. |
| feature_scaling | StandardScaler | Embeddings are standardized before fitting downstream models. |

## D Comparison of Used Atlases

Table 8 summarizes the seven brain atlases used in this study. Among them, five atlases are used during pretraining, while the remaining two atlases are reserved for evaluating generalization to unseen atlases. The Schaefer atlas series (Schaefer100/200/300/400) (Schaefer et al., 2018), one of the most widely used functional parcellation frameworks, was derived from resting-state fMRI data. Its four resolutions, ranging from 100 to 400 regions, provide progressively increasing levels of parcellation granularity while preserving a consistent functional organization, allowing us to evaluate the robustness of the learned representations to variations in the number and granularity of brain regions. The Gordon333 atlas (Gordon et al., 2016) is an independently developed functional atlas with 333 regions, introducing variation in regional definitions beyond the resolution differences within the Schaefer series. For unseen-atlas evaluation, we use Glasser360, a multimodal cortical parcellation with 360 regions, to assess generalization to a distinct regional organization. We also include the Brainnetome Atlas (BNA), a connectivity-based brain atlas constructed from multimodal neuroimaging data that provides a fine-grained parcellation together with detailed anatomical and functional connectivity information. Together, these atlases span diverse parcellation resolutions and regional definitions, enabling a comprehensive evaluation of the atlas-agnostic generalizability of AREION.

**Table 8:** Brain atlases and parcellations used in our experiments.

| Atlas | #ROI | Description |
| --- | --- | --- |
| **Schaefer100** | 100 | Functional parcellation derived from resting-state fMRI and aligned with Yeo 7/17 networks. |
| **Schaefer200** | 200 | Higher-resolution functional parcellation with consistent network organization. |
| **Schaefer300** | 300 | Fine-grained functional parcellation with increased resolution. |
| **Schaefer400** | 400 | High-resolution functional parcellation in the Schaefer atlas series. |
| **Gordon333** | 333 | Functional parcellation derived from resting-state functional connectivity and network organization. |
| **Glasser360** | 360 | Multimodal cortical parcellation with 360 regions. |
| **BNA** | 210 | Connectivity-based brain atlas constructed from multimodal neuroimaging data; 210 cortical regions used in our experiments. |

## E Baselines

To rigorously evaluate the effectiveness of AREION, we compare it with a comprehensive set of baselines, categorized into three groups: traditional machine learning methods, supervised deep learning models, and pretrained fMRI foundation models.

### Traditional Machine Learning Methods

We include a multilayer perceptron (MLP) and random forest (RF) for both classification and regression tasks, along with a support vector machine (SVM) for classification and ridge regression for regression tasks. For SVM and ridge regression, we use the same settings as those used for the downstream adaptation of AREION (see Table 7). For the MLP, we use a three-hidden-layer network with ReLU activations (Nair & Hinton, 2010). The MLP is trained using Adam (Kingma & Ba, 2014) with a learning rate of 10^*−*3^. Additional training and architectural details are provided in Table 9.

**Table 9:** Training and architectural hyperparameters for the MLP and random forest baselines.

| Hyperparameter | Value | Description |
| --- | --- | --- |
| **MLP** |  |  |
| hidden_layer_sizes | (256, 128, 64) | Number and width of hidden layers. |
| activation | ReLU | Hidden-layer activation function. |
| solver | Adam | Weight optimization algorithm. |
| learning_rate_init | $10^{-3}$ | Initial learning rate. |
| alpha | $\{10^{-2}, \dots, 10^4\}$ | 7 log-spaced candidate L2 regularizations. The best value is selected per fold using inner 3-fold cross-validation via GridSearchCV. |
| max_iter | 500 | Maximum number of training iterations. |
| **Random Forest** |  |  |
| n_estimators | 500 | Number of trees in the forest. |
| criterion | Gini / squared error | Split criterion for classification / regression. |
| max_depth | None | Maximum depth of each tree. |
| max_features | sqrt | Uses $\sqrt{d}$ randomly selected features at each split. |

### Supervised Deep Learning Models

We include three supervised deep learning models, including BrainNetCNN (Kawahara et al., 2017), BrainGNN (Li et al., 2021), and BrainNetTF (Kan et al., 2022), covering CNN-, GNN-, and Transformer-based architectures, respectively. All models are trained end-to-end from scratch using supervised learning on the labeled downstream training data without pretraining, and their performance is evaluated on the atlas used for training.

### Pretrained fMRI Foundation Models

For a fair comparison, all foundation models are retrained on our collected data. BrainMass (Yang et al., 2024) uses the same task-specific predictors as AREION, whereas BrainNPT (Hu et al., 2024) and BrainGFM (Wei et al., 2026) are evaluated via end-to-end finetuning following their default protocols. Because BrainMass and BrainNPT support only single-atlas pretraining, we retrain each model separately on every atlas used in the corresponding experiments. Results for Schaefer400 are reported in Table 1, while results for other Schaefer resolutions are provided in Appendix B. BrainGFM, which supports multi-atlas pretraining, is retrained on the same multi-atlas cohort as AREION and evaluated on both Schaefer400 (Table 1) and atlases unseen during pretraining (Table 2). This protocol enables a direct comparison of cross-atlas generalization between BrainGFM and AREION. Figure 6 further compares the prediction performance and model size of the pretrained foundation models on the HCP Schaefer400 sex classification task. AREION achieves the best performance with only 6.4 million parameters, substantially fewer parameters than the other foundation models.

**Figure 6:**
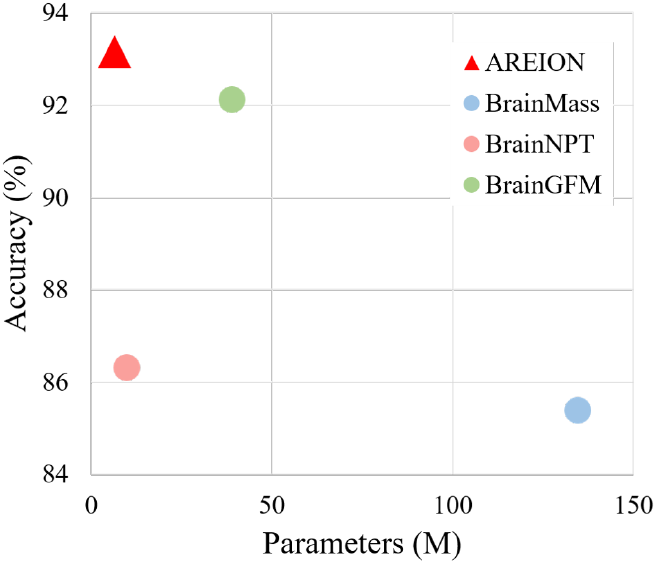
Comparison of performance and model size across different foundation models on the HCP Schaefer400 sex classification task.

## F Interpretability of Learned FC Representations

To assess whether the FC representations learned by AREION capture functional brain organization and its associations with brain development and cognition, we examine regional importance in models predicting age and executive function in the PNC cohort. Regional importance is computed as the region-wise sum of the absolute values of the Haufe-transformed feature weights (Haufe et al., 2014) from each probing model. As shown in Figure 7, cortical regions within the somatomotor network and the ventral attention network exhibit higher importance for age prediction, and regions within the ventral attention network, frontoparietal network, and dorsal attention network show higher importance for the cognition prediction, consistent with previous brain-wide association studies (Dosenbach et al., 2010; Cui et al., 2020; Li et al., 2026; Sripada et al., 2020; Chen et al., 2022; Li et al., 2025), suggesting that AREION captures biologically meaningful variations in its learned FC representations.

**Figure 7:**
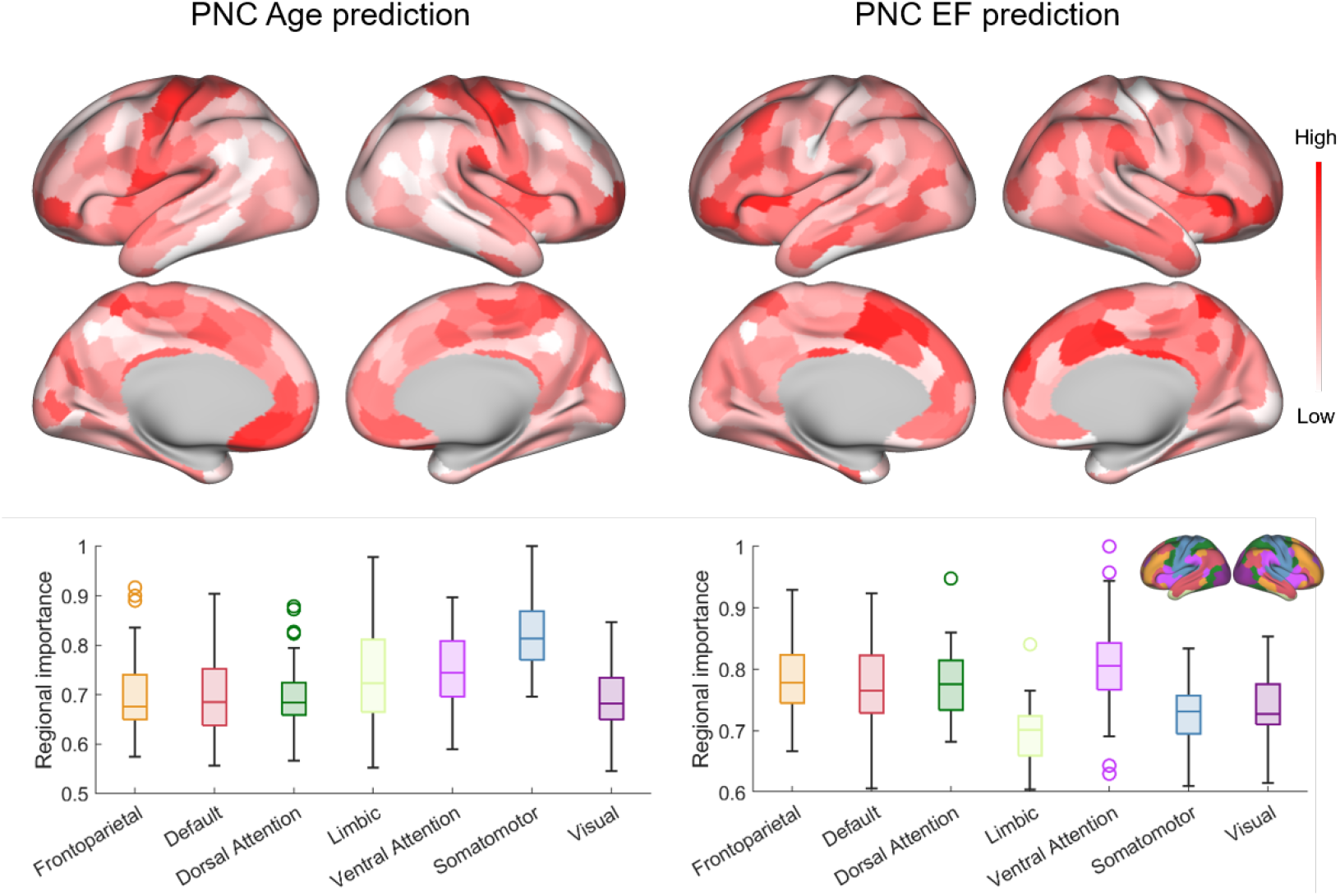
Importance of FC representations for age (left) and executive function (right) prediction across cortical regions (top) and large-scale functional networks (bottom) on the PNC cohort.

